# Pitch of Harmonic Complex Tones: Rate-Place Coding of Resolved Components in Harmonic and Inharmonic Complex Tones in Auditory Midbrain

**DOI:** 10.1101/802827

**Authors:** Yaqing Su, Bertrand Delgutte

## Abstract

Harmonic complex tones (HCT) commonly occurring in speech and music evoke a strong pitch at their fundamental frequency (F0), especially when they contain harmonics individually resolved by the cochlea. When all frequency components of an HCT are shifted by the same amount, the pitch of the resulting inharmonic tone (IHCT) also shifts although the envelope repetition rate is unchanged. A rate-place code whereby resolved harmonics are represented by local maxima in firing rates along the tonotopic axis has been characterized in the auditory nerve and primary auditory cortex, but little is known about intermediate processing stages. We recorded single neuron responses to HCT and IHCT with varying F0 and sound level in the inferior colliculus (IC) of unanesthetized rabbits. Many neurons showed peaks in firing rates when a low-numbered harmonic aligned with the neuron’s characteristic frequency, demonstrating “rate-place” coding. The IC rate-place code was most prevalent for F0>800 Hz, was only moderately dependent on sound level over a 40 dB range, and was not sensitive to stimulus harmonicity. A spectral receptive-field model incorporating broadband inhibition better predicted the neural responses than a purely excitatory model, suggesting an enhancement of the rate-place representation by inhibition. Some IC neurons showed facilitation in response to HCT, similar to cortical “harmonic template neurons” (Feng and Wang 2017), but to a lesser degree. Our findings shed light on the transformation of rate-place coding of resolved harmonics along the auditory pathway, and suggest a gradual emergence of harmonic templates from low to high processing centers.

**Significance statement:** Harmonic complex tones are ubiquitous in speech and music and produce strong pitch percepts in human listeners when they contain frequency components that are individually resolved by the cochlea. Here, we characterize a “rate-place” code for resolved harmonics in the auditory midbrain that is more robust across sound levels than the peripheral rate-place code and insensitive to the harmonic relationships among frequency components. We use a computational model to show that inhibition may play an important role in shaping the rate-place code. We also show that midbrain auditory neurons can demonstrate similar properties as cortical harmonic template neurons. Our study fills a gap in understanding the transformation in neural representations of resolved harmonics along the auditory pathway.

## INTRODUCTION

Pitch is a fundamental attribute of auditory perception that contributes to the comprehension of sounds including speech, music, and animal vocalizations (Bregman, 1994; Plack and Oxenham, 2005b). Most pitch-conveying sounds are harmonic complex tones (HCTs) in which all frequency components are integer multiples of a common fundamental frequency (F0). HCTs that contain low-numbered harmonics individually resolved by cochlear frequency analysis evoke strong pitch sensations at their F0 (see (Plack and Oxenham, 2005a) for review), and provide cues for the perceptual segregation of concurrent sounds (Micheyl and Oxenham, 2010). HCTs containing only high-numbered, unresolved harmonics evoke weaker pitch sensations derived from neural phase locking to the stimulus envelope (Houtsma and Smurzynski, 1990; Shackleton and Carlyon, 1994; Bernstein and Oxenham, 2003). When all harmonics of an HCT with resolved components are shifted in frequency by the same amount, the pitch of the resulting inharmonic complex tone (IHCT) shifts by an amount roughly proportional to the frequency shift although the envelope repetition rate is unchanged (De Boer, 1956; Schouten et al., 1962; Patterson, 1973; Patterson and Wightman, 1976). Although resolved harmonics produce the most salient pitch percepts, the neural mechanisms for extracting pitch from resolved harmonics are poorly understood.

Resolved harmonics may be represented by a “rate-place code” whereby neurons arranged along the tonotopic axis show local maxima in activity when their characteristic frequency (CF) is near a low-numbered harmonic of the stimulus. Such rate-place coding has been described in auditory nerve (AN) fibers of anesthetized cats (Cedolin and Delgutte, 2005; Larsen et al., 2008; Cedolin and Delgutte, 2010), as well as single- and multi-units in the primary auditory cortex (A1) of awake macaques (Schwarz and Tomlinson, 1990; Fishman et al., 2013). While pitch perception is invariant over a wide range of sound level, the AN rate-place code degrades within 20-30 dB above threshold due to saturation of most AN fibers (Cedolin and Delgutte, 2005), although the cortical code appears more robust (Schwarz and Tomlinson, 1990; Fishman et al., 2013). How and where along the ascending pathway level tolerance emerges is unknown. Pitch of harmonic and inharmonic tones may be extracted from tonotopic representations of resolved harmonics using harmonic templates (Goldstein, 1973; Wightman, 1973; Terhardt, 1974; Cohen et al., 1995; Shamma and Klein, 2000). However, there is little physiological evidence for harmonic templates, except for a report of “harmonic template neurons” (HTNs) in the auditory cortex of marmoset monkeys (Feng and Wang, 2017). These neurons show firing rate facilitation at a particular F0 of HCT compared to a pure tone at CF, and are sensitive to stimulus harmonicity.

The inferior colliculus (IC) is a logical site to investigate how rate-place coding transforms along the auditory pathway, and possibly shed light on the emergence of harmonic templates. Its tonotopically organized central nucleus receives convergent excitatory and inhibitory inputs from most brainstem auditory nuclei (Adams, 1979; Malmierca et al., 2005), as well as inputs from within IC (Saldana and Merchan, 1992). A handful of IC studies that used HCT stimuli containing many harmonics (Sinex et al., 2002; Sinex and Li, 2007; Shackleton et al., 2009; Schnupp et al., 2015; Peng et al., 2018) focused on the coding of unresolved harmonics, and mostly tested low F0s within the range of human voice, which are not likely to be resolved in experimental animals.

Here, we characterize rate-place coding of resolved harmonics in the IC of unanesthetized rabbits by measuring single-unit responses to HCT and IHCT with varying F0 and sound level. Spectral receptive field models were fit to neural responses to evaluate the role of inhibition in shaping the IC rate-place code. We also compared responses to HCT and pure tone stimuli to assess whether IC neurons exhibit properties of cortical HTNs (Feng and Wang, 2017). We find that IC neurons can represent resolved harmonics over a wide range of F0 via a rate-place code that is robust across sound levels and not sensitive to harmonicity.

## METHODS

Four female and one male adult Dutch-belted rabbits were used for the experiments. All animal procedures we approved by the Animal Care Committee of Massachusetts Eye & Ear.

### Surgical preparation

Surgical procedures were adapted from Kuwada and colleagues (Kuwada et al., 1987) and have been described in previous publications from our laboratory (Devore and Delgutte, 2010; Day et al., 2012). Each rabbit underwent two aseptic surgeries before the first electrophysiological recording session: a head bar and cylinder implantation, and a craniotomy. In both surgeries, anesthesia was induced with xylazine (6 mg/kg) followed by ketamine (35-44 mg/kg), and maintained by either of the two methods: 1) injection of 1/3 initial dose of xylazine/ketamine mix when the animal showed a withdrawal reflex, 2) facemask delivery of isoflurane gas mixed with oxygen (0.8 l/min, isoflurane concentration gradually increased to 2.5%).

#### Head bar and cylinder implantation

A brass head bar and a stainless steel cylinder were affixed to the skull with stainless steel screws and dental acrylic. At the end of the surgery, the exposed skull was covered with topical antibiotic ointment (Bacitracin) and the cylinder filled with vinyl polysiloxane (Reprosil). This covering procedure was also applied after the craniotomy and at the end of every electrophysiological recording session.

#### Craniotomy

Once fully recovered from the initial surgery, the rabbit was trained to sit still in the experimental apparatus with head attached over the following 8-10 days. Then, a small craniotomy (2-3 mm diameter) was then made within the cylinder at 10.5 mm posterior and 3 mm lateral to bregma to allow access to the IC. Immediately after the craniotomy, custom ear inserts were cast using Reprosil. Occasionally over the period of recording sessions, the craniotomy was enlarged or moved to the contralateral side using the same procedure.

### Single-unit recording

Each recording session lasted 1.5-2.5 h, during which the rabbit was strapped in a spandex sleeve with its head fixed via the brass bar in a double-walled, electrically shielded and sound-proof chamber. At the beginning of each session, the acoustic pressure inside the ear canal in response to a broadband chirp stimulus was measured with a probe-tube microphone (Etymotic ER-7C) to calibrate the acoustic assembly. An inverse digital filter was then created from the calibration over 0.05-18 kHz. The animal was monitored via a close-circuit video throughout the session, and the recording session was terminated if the animal showed signs of distress or moved excessively.

The majority of single neuron recordings were made with polyimide-insulated platinum-iridium linear microelectrode arrays (MicroProbes) with 4-6 contacts spaced 150 μm apart, 0.2-1 MΩ impedance each. Some early recordings were made using epoxy-insulated tungsten electrodes (A-M Systems) with 2-4 MΩ impedance. During recording, the electrode was advanced through the occipital cortex towards the IC by a remote-controlled hydraulic micropositioner (David Kopf Instruments Model 650). The IC was identified by audio-visual cues of entrainment to a search stimulus consisting of 200-ms broadband noise bursts presented diotically at 60 dB SPL. The signals recorded from the microelectrode array were first amplified and bandpass filtered from 300 to 5000 Hz (Plexon, PBX2), then sampled at 100 kHz (National Instruments, PXI-6123). Spike times were identified by crossing of a manually-set voltage threshold and recorded for later analysis. The signal recorded from tungsten electrodes was amplified (Axon Instruments, Axoprobe-1A) and filtered (Ithaco 1201) then processed in the same way. Isolation of a single unit was determined based on the stable shape and amplitude of the spike waveform and absence of interspike intervals < 1 ms.

### Stimuli

Acoustic stimuli were first created in MATLAB (The MathWorks) and passed through the digital filter created from the acoustic calibration to equalize the frequency response of the acoustic assembly. The filtered signals were then converted to analog signals by a 24-bit D/A converter (National Instruments, PXI-4461) and delivered to the animal ear by a pair of speakers (Beyer-Dynamic, DT48) via plastic tubes fitted through the ear inserts. Once a neuron was isolated, we characterized its frequency tuning with pure tones and then measured responses to complex tones.

#### Pure tone frequency tuning

In half of the neurons, we measured the frequency response area (FRA) to characterize pure-tone tuning. Tone bursts (100-ms on, 100-ms off) varying in frequency from 200 Hz to 18 kHz (0.25 octave step or finer) and in level from 5 dB SPL to 70 dB SPL were presented in random order and each was repeated 3 times. The evoked firing rate was measured for each tone, and plotted as a heat map on the log frequency vs. intensity plane. The heat map was then interpolated 10x. Contours on the interpolated map were identified using the MATLAB image processing toolbox. The characteristic frequency (CF) was defined as the frequency corresponding to the lowest sound level on the longest contour (see Figure 4 for examples).

Before the FRA measurement was implemented, pure-tone frequency tuning was characterized with either an automatic threshold tracking algorithm or an iso-level method. In the tracking method (Kiang and Moxon, 1974), tone bursts (125-ms on, 125-ms off) were presented from high (18 kHz) to low (50 Hz) frequencies in 0.1 octave steps, and an automatic algorithm determined the threshold level at each frequency. The CF was defined as the frequency yielding the lowest threshold. This method often failed for neurons in which the FRA was irregular in shape or showed inhibitory zones. When the tracking method failed, we used an iso-level method in which tone bursts varying in frequency from 0.5 to 18 kHz (200-ms on, 300-ms off, 0.1 octave steps) were presented in random order at a fixed level (approximately 10 dB above the threshold to broadband noise) with 3 repetitions. The best frequency (BF) was defined as the frequency that evoked the highest firing rate during tone presentation.

Most pure tone responses were measured for monaural stimulation of the contralateral ear. In rare cases when a neuron responded more strongly to ipsilateral sounds, frequency tuning was characterized for monaural stimulation of the ipsilateral ear. For brevity, we will refer to both the characteristic frequency measured from the FRA or the tracking method and the best frequency measured by the iso-level method as “CF” in this report.

#### Harmonic complex tones

Rate-place coding of resolved harmonics by single neurons was tested using a harmonic complex tone paradigm adapted from previous studies in the auditory nerve (Cedolin and Delgutte, 2005; Larsen et al., 2008) and auditory cortex (Fishman et al., 2013). This paradigm was designed to optimize the chance of observing responses to resolved harmonics by selecting the most appropriate range of F0s based on each neuron’s CF. Each HCT consisted of equal-amplitude harmonics in cosine phase, ranging from the fundamental (F0) to the lowest of either the 12^th^ harmonic or 18 kHz, where the frequency response of our acoustic system showed a sharp drop-off. The power spectrum and temporal waveform of a harmonic complex with F0 equal to 500 Hz are shown in Figure 1 (2^nd^ row, orange).

**Figure 1.**
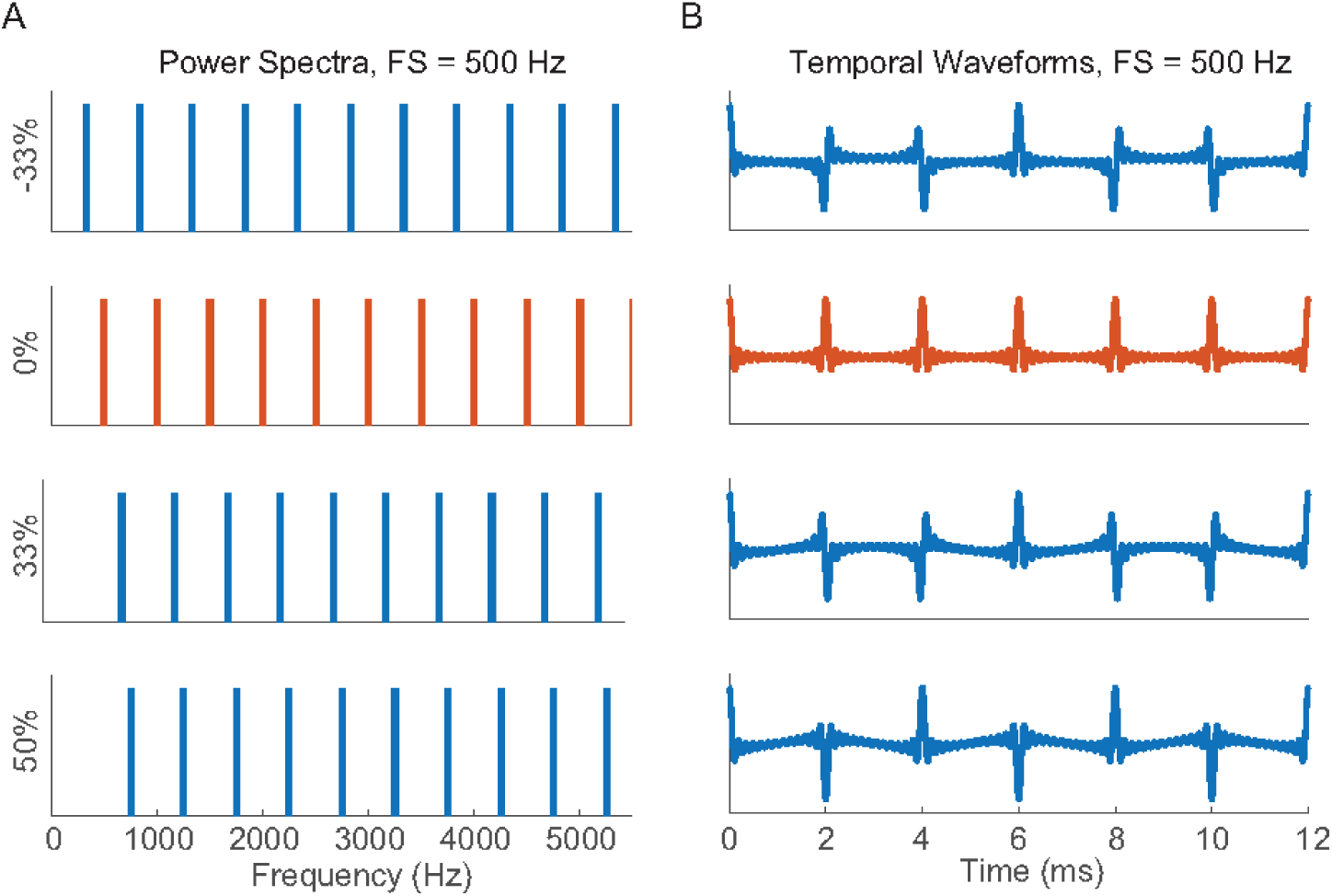
Harmonic and inharmonic complex tones. **(A)** Power spectra of harmonic complex tone (0% shift, orange) and inharmonic complex tones (non-zero shift, blue) with the same frequency spacing FS=500 Hz. Y labels indicate amounts of frequency shift from the harmonic complex tone as percentages of FS. For the harmonic tone, FS is equivalent to F0. **(B)** Temporal waveforms of the corresponding complex tones in **A**.

For each neuron, the F0 of HCTs was varied so that the “neural harmonic number”, NH=CF/F0, varied from 0.5 to 5.5 or higher in linear steps of 1/6. When NH is a small integer (Figure 2A), the NH^th^ component of the complex tone coincides with the neuron’s CF, therefore the stimulus should evoke a high firing rate is this harmonic is resolved. When NH is an integer + 1/2 (Figure 2C), the neuron’s CF falls half way between two harmonics, so that a low firing rate is expected if the harmonics are resolved. As NH increases (Figure 2D), F0 decreases so that the spacing between adjacent harmonics (i.e. F0) becomes smaller than the bandwidth of the neuron’s FRA, so the firing rate should no longer depend on F0. In this way, HCTs with low-number harmonics (large F0) are expected to be resolved by the neuron, while HCTs with high-number harmonic (small F0) should be unresolved.

**Figure 2.**
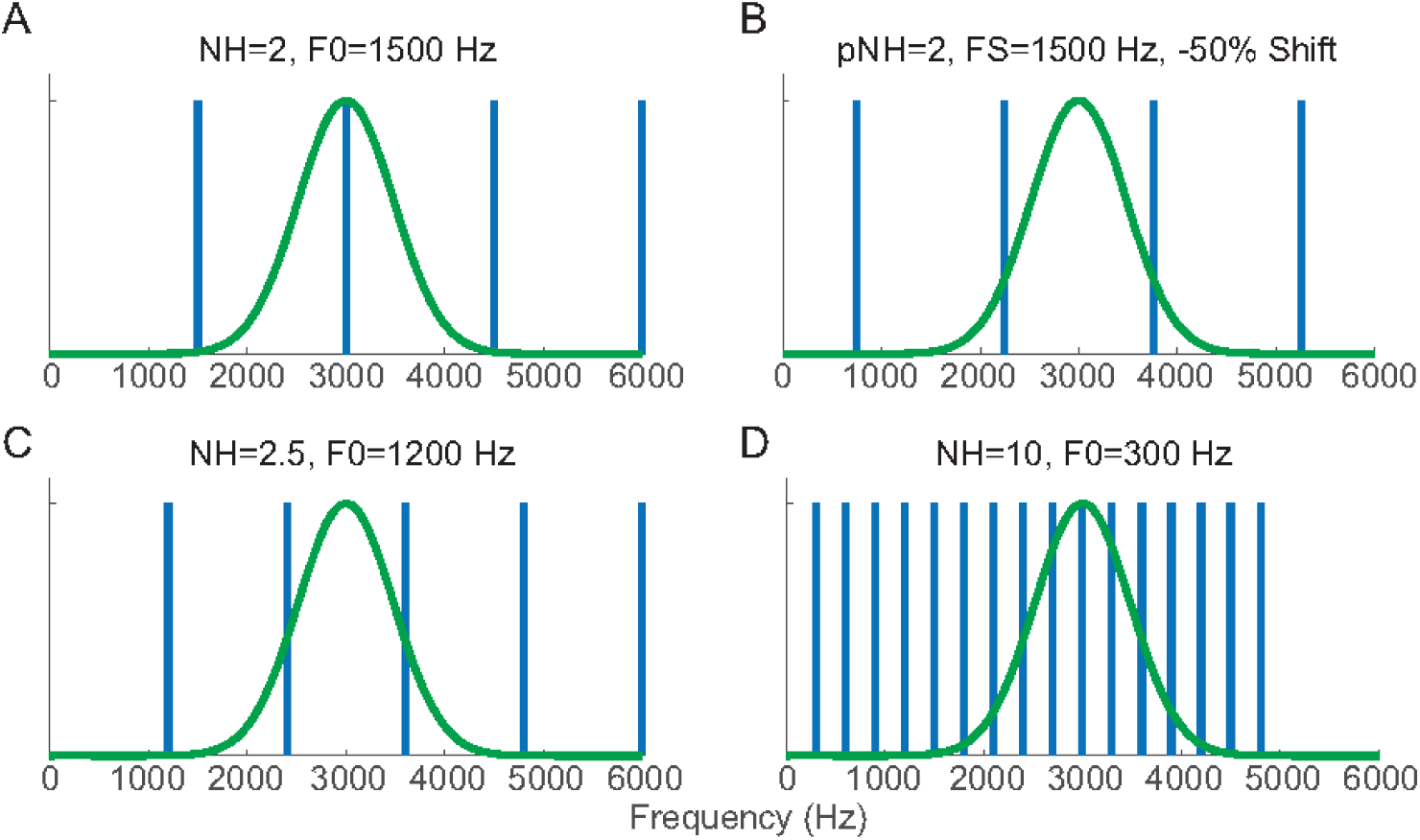
Schematics for testing rate-place coding in a hypothetical neuron with CF=3000 Hz (green curves). Vertical bars represent power spectra of the corresponding complex tone. **(A)** NH=2 (F0=1500 Hz), the second harmonic of the corresponding HCT coincides with the neuron’s CF. **(B)** For an inharmonic complex tone with the same frequency spacing as in A but −50% shift (750 Hz to lower frequency), the neuron’s CF is between two adjacent components. **(C)** NH=2.5 (F0=1200 Hz), CF between two adjacent harmonics. **(D)** NH=10, (F0=300 Hz), the neuron’s auditory filter encompasses many harmonics.

HCTs with each F0 were presented at three different sound levels: low (<=30 dB SPL/harmonic), medium (31-55 dB SPL/harmonic) and high (55-70 dB SPL/harmonic). Sound level was specified for each harmonic rather than overall so that the amplitude of individual harmonics would stay the same when F0 is varied. When the stimulus contains 12 harmonics, the overall SPL is about 11 dB higher than the level per harmonic. The three sound levels used in each neuron were originally chosen in 15 dB increments, but this was later increased to 20 dB increments to test a wider range. In each measurement, HCTs with different F0s and sound levels were randomly interleaved for 10 total repetitions. Each complex tone was presented diotically for 200 ms with a 10-ms raised-cosine ramp at onset and offset, and followed by a 300-ms silent (off) period.

For a subset of neurons, HCT stimuli were interleaved with pure tones at frequencies such that the ratio CF/F ranged from 0.5 to 1.5 in steps of 1/6, with the same duration (200 ms), inter-stimulus interval (300 ms), and per-harmonic sound levels as the HCTs.

#### Inharmonic complex tones

In a subset of neurons, we measured responses to inharmonic complex tones (IHCT) that were generated by shifting all frequency components in a HCT by a fixed proportion of its F0. For an inharmonic tone, the frequency spacing (FS) between adjacent components is always equal to the F0 of the original HCT, and the temporal envelope repetition rate is 1/FS (Figure 1). For the majority of neurons, inharmonic tones were generated with shifts of ±1/3, ±1/6, 0 (harmonic) and ½ of the original F0. In a few earlier measurements, frequency shifts of 0, ±1/10, ±1/4 and ½ were used.

A pseudo-harmonic number (pNH) was defined for each IHCT as pNH=CF/FS. For each neuron and each value of frequency shift, pNH was varied from 0.5 to 6.5 as NH for the harmonic tones. Because of the frequency shift, an IHCT with integer pNH no longer has a component at the neuron’s CF (Figure 2B). Instead, for the *N^th^*component of an IHCT with shift *s* to align with the neuron’s CF, the tone should satisfy

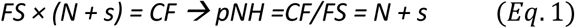

Therefore, if a neuron’s response is dependent on the presence of resolved components near the CF regardless of harmonicity, it would show peaks in firing rates when *pNH = N* (a small integer) *+ s* (percent shift relative to FS).

Due to the large number of stimulus conditions tested (36 FS values × 6 frequency shifts), IHCTs were only presented at 30 dB SPL per component.

### Data analysis

For all HCT and IHCT paradigms, we computed the average firing rate over the stimulus duration as a function of (pseudo) harmonic number. The response in the first 10 ms was excluded to eliminate the onset response. The neuron’s background firing rate was calculated as the average firing rate during the last 200 ms of the 300 ms inter-stimulus-interval averaged over across all the complex tone stimuli. Assuming that there exists a population of IC neurons having the same response properties as the recorded neuron, except for their position along the tonotopic axis (mapped to their CF), a plot of firing rate against neural harmonic number (or F0) should resemble the response of the hypothetical neural population to a HCT with a specific F0 as a function of CF (Larsen et al., 2008). Therefore, we refer to the patterns of firing rate against neural harmonic number as “rate-place profiles”. This terminology is consistent with previous studies of the auditory nerve and cortex (Cedolin and Delgutte, 2005; Fishman et al., 2013).

#### Fourier-based analysis of rate-place profiles

The rate-place profile of a neuron demonstrating rate-place coding should have peaks at small integer harmonic numbers and troughs in between small integers, producing a periodic pattern with periodicity of 1 (in units of harmonic number NH). Following Fishman et al. (2013), we harnessed this periodicity by using a Fourier-based analysis to characterize rate-place coding in our neurons (see Figure 4B-D for example).

The discrete Fourier transform (DFT) of the rate-place profile was first computed and normalized by half the average firing rate across the entire profile so that the DFT amplitude fell in the range of [0, 2]. The component at zero frequency (DC) was then set to zero to simplify the following analysis. The “harmonic modulation depth” was defined as the amplitude of the Fourier component at 1 cycle/harmonic number. With a periodicity of one cycle/NH, a fully modulated sinusoidal profile would have a harmonic modulation depth of 1, while a profile consisting of impulses at integer values of NH would have a depth of 2. The statistical significance of the harmonic modulation depth was determined by a permutation test (10,000 permutations), where data points on the rate-place profile were randomly shuffled across NH and the harmonic modulation depth was computed for each permutation. The neuron was identified as showing rate-place coding if the original modulation depth exceeded the 95 percentile of the permutation values.

##### CF adjustment

We observed that the periodicity of rate-place profiles was sometimes not exactly at, but still close to, one cycle/harmonic number. This was possibly due to inaccuracy in CF measurement or to differences in frequency selectivity between pure tones and HCT as a result of nonlinear processing. Such deviations from the expected periodicity could accumulate and have a pronounced effect on rate profiles at large harmonic numbers. We adapted a method from Fishman et al. (2013) to adjust for these deviations and obtain a revised estimate of CF that better describes responses to HCT. Specifically, we estimated the location of the maximum of the Fourier spectrum by fitting a parabola to the Fourier spectrum at the peak location and the two neighboring points, and then computing the location of the peak, φ, of the fitted parabola. This procedure was done for each of the three sound levels separately, and the final φ value was the average across all levels at which the harmonic modulation depth was statistically significant. The adjusted CF was defined as CF_adjusted_=φ×CF_measured_ because the adjustment is equivalent to scaling the DFT axis by a factor φ so that the parabola peak occurs exactly at one cycle/NH. All NH values corresponding to the tested F0s were adjusted as well using the formula NH_adjusted_ = CF_adjusted_/F0= φ×NH_nominal_. The harmonic modulation depth was also redefined as the amplitude of the scaled Fourier spectrum at one cycle/NH.

##### Identification of resolved harmonics

Peaks in the rate-place profile corresponding to resolved harmonics were identified by applying the Fourier analysis recursively. When a significant spectral peak at one cycle/NH was identified in the Fourier spectrum of the rate-place profile, data points for the first peak (NH from 0.5 to 1.5) were removed from the profile. The same procedure was then applied to the remainder of the rate-place profile, and repeated until the harmonic modulation depth became insignificant. For a profile with N peaks identified by this procedure, the total number of resolved harmonics is N+1, because a profile must have at least 2 peaks to show periodicity. Thus a neuron that would only resolve the fundamental but not the second harmonic would not be considered to exhibit rate-place coding based on our criteria.

#### Quantification of neural coding strength

To characterize neural sensitivity to F0 independently of whether the rate-place profile showed resolved harmonics or not, we computed the signal-to-total variance ratio (STVR) (Hancock et al., 2010; Hancock et al., 2012). STVR is an ANOVA-based metric derived from the spike counts on each stimulus trial that represents the ratio of the variance in firing rates attributable to their dependence on F0 to the total variance, including variability across multiple presentations of the same stimulus. STVR=1 implies perfectly reliable sensitivity to F0, i.e. all the response variability can be attributed to the change in F0, and 0 implies no sensitivity (flat rate profile).

#### Spectral receptive fields models

For neurons demonstrating resolved harmonics in HCT responses, we used linear spectral receptive field models to predict the rate-pace profile from the stimulus spectra at each F0 (Figure 3). Two spectral receptive field models were tested—a simple Gaussian function:

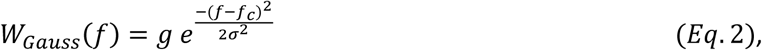

and a Difference of Gaussians (DoG) function to simulate the interaction of excitation and inhibition:

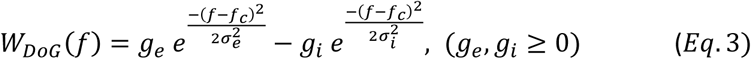

**Figure 3.**
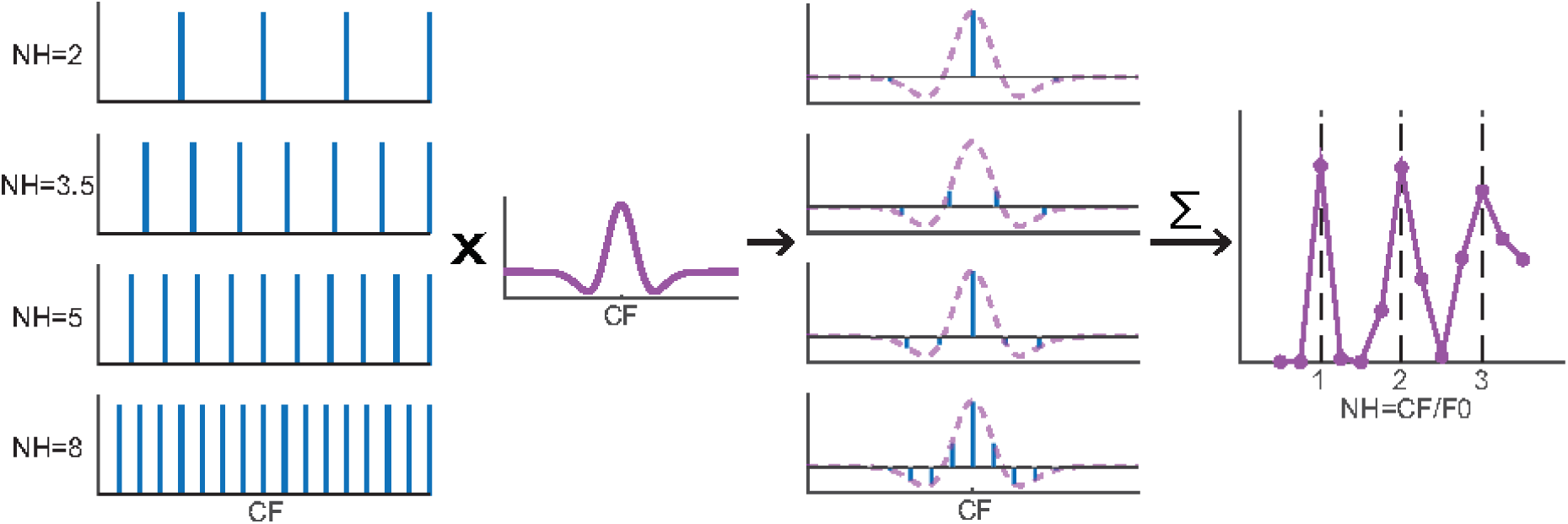
DoG spectral weighting model diagram. **Column 1:** Power spectra of HCTs at different NHs. **Column 2:** A DoG weighting functions centered at the neuron’s CF. **Column 3:** Weighted power spectra by multiplying the HCT spectra with the weighting function (purple dashed lines). Horizontal lines in each panel indicate zero amplitude. **Column 4:** Modeled rate-place profile of the neuron. Each point is obtained by summing the weighted power spectrum in Column 3 of an HCT at the corresponding F0, or equivalently, the convolution of the power spectrum and the weighting function at zero shift.

**Figure 4.**
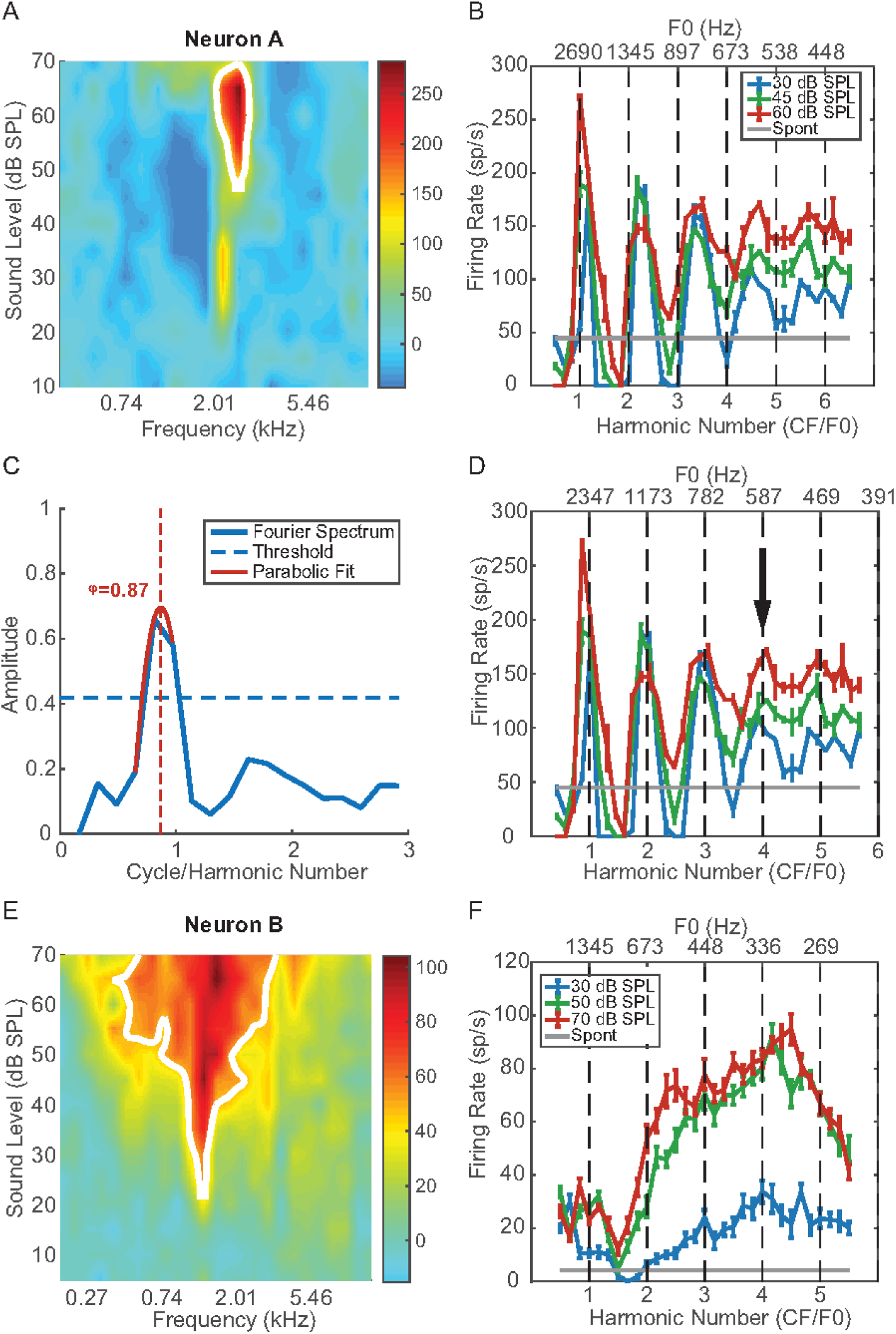
Example neural responses to harmonic complex tones. **(A)** Pure tone frequency response area of Neuron A. Colors indicate the neuron’s firing rate minus the spontaneous rate. **(B)** Rate-place profile of Neuron A before the CF adjustment (see Methods). **(C)** Fourier transform of the rate-place profile at 30 dB SPL in **B**. Horizontal dashed line indicates the 95 percentile of the permutation test, vertical dashed line indicates the location of the “true” spectrum peak, i.e. the adjustment factor φ, inferred from the parabolic fit (red solid line). **(D)** Rate-place profiles of Neuron A after adjustment. The arrow indicates the neuron’s lowest resolved F0. **(E)** Pure tone FRA of Neuron B, CF=1.35 kHz. **(F)** Rate-place profile of Neuron B.

In both functions, the amplitude *g* has a unit of neural firing rates in spike/s. The center frequency *f_c_* is expected to correspond to the neuron’s CF, and the standard deviation σ specifies bandwidth for the corresponding Gaussian filter. For both models, the firing rate *r* is computed from the stimulus spectrum *S(f)* by the equation:

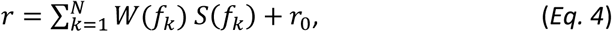

where the *f_k_* represent the frequency components of the stimulus and *r_0_* is the spontaneous firing rate. Parameters of the two models were fitted separately by minimizing the city-block distance (sum of absolute distances across F0s) between the model prediction and the neuron’s rate profile using the MATLAB function *fmincon*. The fitted rate profiles were half-wave rectified, i.e. any negative predicted firing rates were set to 0 in the model output. Goodness-of-fit of each model was assessed using adjusted R-squared (*n*—number of data points; p—number of parameters) to take into account the different number of parameters in the two models (Theil, 1961):

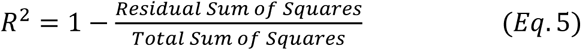

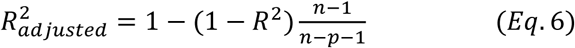

We also compared the goodness of fit the two models using single-sided F tests, for which the null hypothesis was that the two models fit equally well, and the alternative hypothesis that the DoG fit better than the Gaussian model.

The Gaussian function has a purely excitatory (g>0) or inhibitory (g<0) band centered at *f_c_*, while the DoG has a more complicated morphology depending on the interaction between the excitatory and the inhibitory components. For most neurons, the inhibitory component of the best fitting DoG model had a smaller amplitude and wider bandwidth than the excitatory component (g_i_<g_e_, σ_i_> σ_e_), resulting in a narrower excitatory center band flanked by two symmetrical inhibitory sidebands as illustrated in Figure 3. In cases where σ_i_< σ_e_, the receptive field has a broad excitatory band with a notch in its center.

#### Experimental design and statistical analysis

Each neuron’s responses to different stimulus conditions (F0s, sound levels or frequency shifts of inharmonic complex tones) were obtained using randomly interleaved presentations to minimize the effect of possible fluctuations in overall neural responsiveness. Whenever possible, we used nonparametric statistical tests to compare neural response metrics (e.g. the STVR) between stimulus conditions across the neuronal population. When comparing two conditions, we used the Wilcoxon signed rank test (for related variables) or rank-sum test (for independent variables), and the Kolmogorov-Smirnov test (KS test) for comparing the distributions. For three or more conditions, we used the Friedman test (Friedman, 1937) to compare across all conditions, and obtained pairwise comparison by applying multiple comparison (with Bonferroni correction) to the Friedman test statistics. We used the chi-squared test for comparing distributions of discrete data such as the number of resolved harmonics among different sound levels. Significance of the correlation between two quantities was determined by the Kendall tau test (Kendall, 1948). For all tests, p<0.05 was considered as statistically significant. All statistical tests were performed using the MATLAB statistics toolbox.

## RESULTS

We recorded responses to HCT stimuli from 252 IC neurons in five unanesthetized rabbits. 194 of these neurons had an identifiable CF so that the range of F0 could be set based on the desired range of neural harmonic numbers (0.5 to 6.5) to test for rate-place coding of resolved harmonics. Responses of these 194 neurons to HCTs were studied as a function of F0 and sound level. The remaining 58 neurons that did not have a clearly identifiable CF were used to study responses to HCT with unresolved harmonics as reported in a companion paper (Su and Delgutte, 2019). These 58 neurons are not included in the present data set. Among the 194 neurons included in the present data set, 25 were also tested with inharmonic complex tones, and 37 were tested with pure tones interleaved with the HCT to determine whether they behave like cortical harmonic template neurons (Feng and Wang, 2017). Pure tone CFs of the neurons ranged from 0.4 to 24.3 kHz, with a median of 4.6 kHz.

### Rate-place coding of resolved harmonics in HCT

Figure 4 shows the pure-tone frequency response areas (FRA) and HCT rate profiles of two IC neurons. Neuron A (Figure 4A-D) had a sharply-tuned, nearly “I” shaped FRA with two excitatory zones. The contour-based algorithm (see Method) identified 2.69 kHz (the tip of the high-threshold zone) as the CF (Figure 4A), and this value was used to set the range of F0s of the HCT stimuli from 414 Hz (NH=6.5) to 5380 Hz (NH=0.5). The rate-place profiles (Figure 4B) showed peaks near the first four integer harmonic numbers at all three stimulus levels tested, but were “stretched” so that the inter-peak intervals were slightly larger than 1 NH. Figure 4C shows the Fourier transform of the neuron’s rate-place profile at 30 dB SPL. The peak amplitude, i.e. the harmonic modulation depth, was 0.66, which exceeded the 95% percentile of the permutation test (horizontal dashed line). Using parabolic interpolation, the peak in the spectrum was identified at φ = 0.87 cycle/NH (vertical dashed line) rather than the predicted 1 cycle/NH. Therefore, the CF was adjusted to 0.87×2.69 = 2.34 kHz. The adjusted CF corresponds to the low-threshold zone of the FRA, which was not identified as the CF by the automatic algorithm because its contour had a shorter length than the high-threshold zone (white line in Figure 4A). After CF adjustment, peaks in the rate profiles were aligned at integer NHs (Figure 4D). For neurons showing peaks in firing rate at resolved harmonics in their rate-profile, like Neuron A, we define the “lowest resolved F0” as the F0 corresponding to the largest integer NH that yielded a significant peak the rate profile according to the DFT-based algorithm (see Methods). The lowest resolved F0 for Neuron A was 587 Hz (4^th^ harmonic, arrow on Figure 4D) for all three sound levels.

Although Neuron B had a “V” shaped pure tone FRA with a clear CF at 1.35 kHz (Figure 4E), unlike Neuron A, its rate profile failed to show peaks in firing rate near integer harmonic numbers (Figure 4F), and the peak in the spectrum of the rate profile lay below the 95% percentile of the shuffled values (not shown). Such “non-place coding” neurons were a majority in our sample. In total, 80/194 neurons (41%) demonstrated rate-place coding of resolved harmonics, while the remaining 59% showed no evidence of rate-place coding. Our experimental design only measured responses to HCT in neurons with a clearly identifiable CF in response to pure tones. It is unclear how this selection may have affected the estimate of the proportion of rate-place coding neurons because, in highly nonlinear neurons as found in IC, it is theoretically possible that a neuron without a pure tone CF would still show harmonically related peaks in firing rate in response to HCTs. Nevertheless, it seems likely that the proportion of rate-place coding neurons was overestimated overall.

Rate-place coding of resolved harmonic was observed in IC neurons across a wide range of CFs. Figure 5A shows the distribution of adjusted CF for place coding (dark green) and unadjusted CF for non-place-coding (light green) neurons. The distributions for both groups extended from a few hundred Hz to above 10 kHz with more neurons at higher CFs. However, the median CF was significantly larger for place coding neurons (6998 Hz) than for non-coding neurons (3707 Hz) (p=7.25×10^-5^, Wilcoxon rank-sum test).

**Figure 5.**
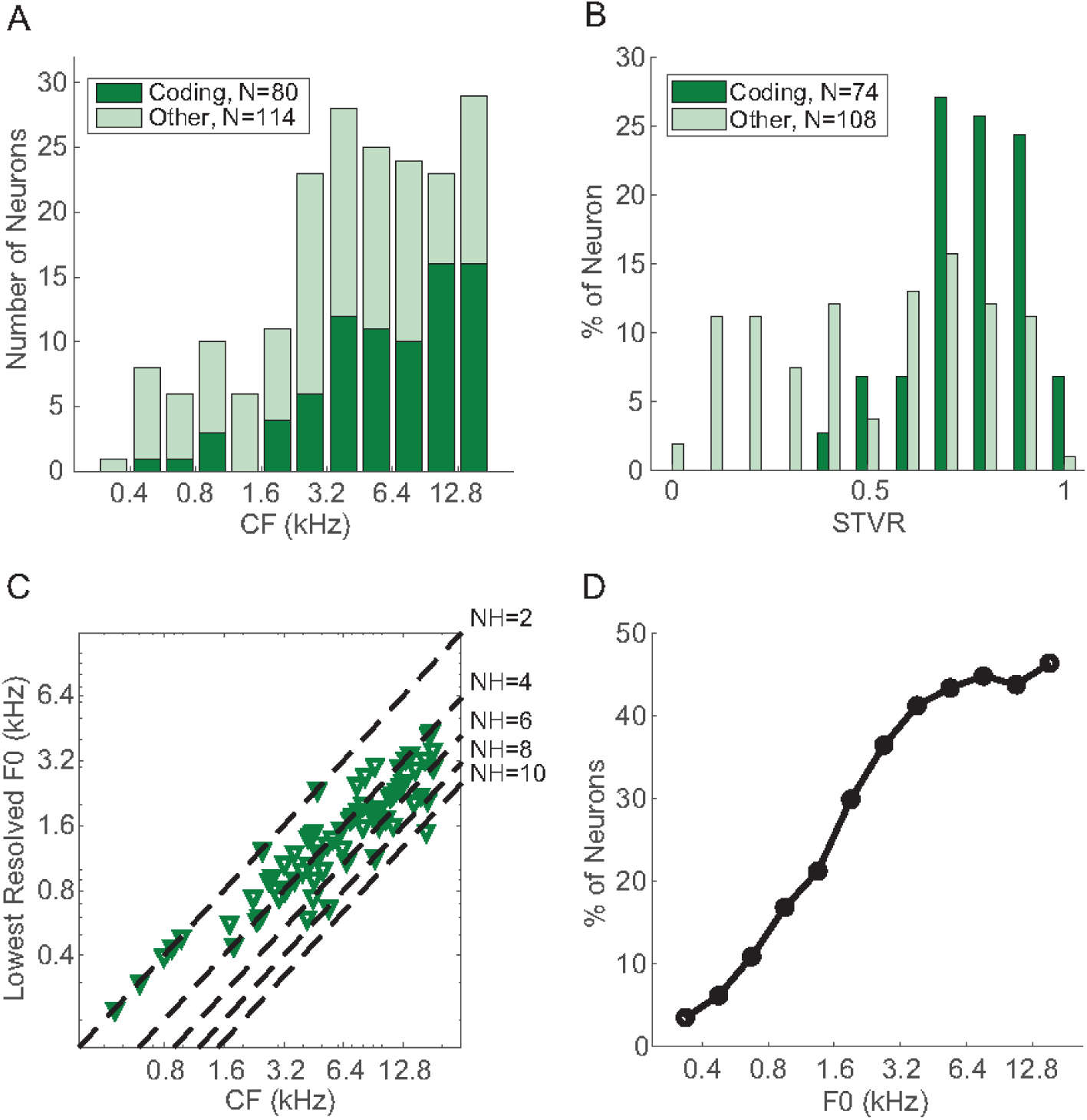
Frequency distribution of IC rate-place code. **(A)** Distribution of CF for rate-place coding neurons (dark green, bottom) and non-rate-place coding neurons (light green, top). **(B)** Distribution of STVR for coding (dark green) and non-coding (light green) neurons. Neurons tested with less than five total stimulus repetitions were excluded. **(C)** Relationship between the lowest resolved F0 and CF of individual rate-place coding neurons (N=80). Dashed lines indicate NH=2, 4, 6, 8, 10. **(D)** Percentage of neurons that were able to resolve F0 in different frequency bands.

We used an ANOVA-based metric, the “signal to total variance” ratio (STVR, see Methods) to quantify the strength of F0 coding in both groups of neurons independently of the shape of their rate-place profiles. The distribution of F0 STVRs (Figure 5B) was skewed towards higher values (close to 1) in place coding neurons, but was relatively uniform for non-coding neurons, and the median STVR was higher for place-coding neurons than for non-coding neurons (0.80 vs. 0.58, p<10^-9^, single sided Wilcoxon rank sum test test). A higher STVR implies that a greater amount of the variance in firing rates can be explained by the variation in F0 as opposed to intrinsic variability in neural firing. The greater ability of place-coding neurons to reliably encode F0 justifies our focus on this group of neurons in the remainder of this paper.

For each place-coding neuron, the lowest resolved F0 over all three sound levels (e.g. 587 Hz for Neuron A) is plotted as a function of adjusted CF in Figure 5C. Neurons with CF<1000 Hz were only able to resolve the first two harmonics, and the lowest resolved F0 found among all neurons was 300 Hz, corresponding to the 2^nd^ harmonic of a neuron with CF=600 Hz. Neurons with higher CFs were able to resolve more harmonics (typically 3-5, but up to 11 in one case), but the lowest resolved F0 remained above 400 Hz. To estimate the availability of the rate-place code as a function of F0, we computed the proportion of neurons that were able to resolve an F0 within a ½-octave frequency band relative to the total number of neurons tested with HCTs in that band (including non-place-coding neurons). As shown in Figure 5D, the proportion monotonically increases with F0, and reaches a plateau of approximately 40% for F0 of 3200 Hz or higher. Thus, rate-place coding is more effective at high F0s compared to low F0s, consistent with the improvement in cochlear frequency selectivity (Q10) with increasing CF (Borg et al., 1988).

### Moderate dependence of IC rate-place coding on sound level

For human listeners, pitch perception of HCT is robust across a wide range of sound levels (Zheng and Brette, 2017). In contrast, rate-place coding in the auditory nerve degrades at levels 20-30 dB above threshold due to spike rate saturation (Cedolin and Delgutte, 2005). In our neuronal sample, many IC neurons demonstrated strong rate-place coding at high sound levels (>55 dB and up to 70 dB SPL per harmonic) that was similar to their response at lower sound levels in terms of harmonic resolvability. To characterize the dependence of IC rate-place coding on sound level, we used three different measures. First, we compared the “harmonic modulation depth” of the spectrum of the rate-place profile at low (≤30 dB per harmonic), medium (31-55 dB) and high (>55 dB) SPLs in neurons showing rate-place coding for at least one sound level (Figure 6A). In general, the harmonic modulation depth tended to decrease slightly with increasing sound level, with medians of 0.45, 0.41, and 0.35 for low, medium and high SPL, respectively (compare to 0.66 for Neuron A in Figure 4 at low level). Differences in median harmonic modulation depths were significant across all three sound levels (p=3.5×10^-6^, Friedman test, N=79), and also between low vs. medium (p=0.026, multiple comparison on the Friedman test statistic with Bonferroni correction) and low vs. high (p=1.6×10^-6^) sound levels. The difference between medium and high sound levels was close to significance (p=0.051).

**Figure 6.**
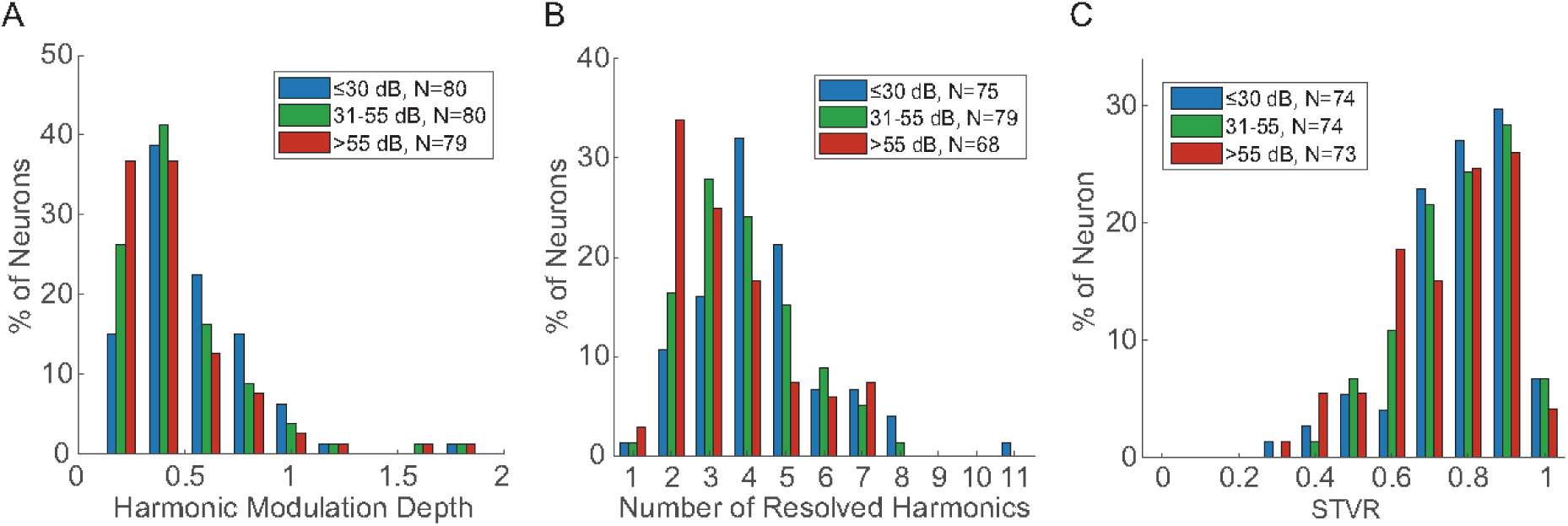
Level dependence of IC rate-place code. **(A)** Distribution of harmonic modulation depth for all neurons demonstrating rate-place coding in at least one sound level. **(B)** Distribution of total number of resolved harmonics at three sound levels. Only neurons showing resolved harmonics at the corresponding sound level are included. **(C)** Distributions of STVR at three sound levels in place-coding neurons.

Figure 6B shows the distributions of the number of resolved harmonics in the three sound level ranges for rate-place coding neurons. The distributions spanned a similar range for all three sound levels, but their centroids shifted slightly towards lower numbers as sound level increased, indicating a reduction in the effective frequency range of rate-place coding at higher sound levels. The difference among the three distributions was statistically significant (p=0.037, chi-square test). (The numbers of neurons contributing to the histogram were smaller than the total number of rate-place coding neurons, because some neurons only showed rate-place coding at one or two of the sound levels.)

Figure 6C compares the distribution of F0 STVR among place-coding neurons across sound levels. Median STVR values were 0.80, 0.81 and 0.77 for low, medium and high SPL respectively. Differences in median STVRs were significant across all three sound levels (p=8.9×10^-4^, Friedman test, N=73), but pairwise comparison revealed only a significant difference between low and high sound levels (p=0.41 for low vs. mid, p=0.076 for mid vs. high, and p=5.9×10^-4^ for low vs. high, multiple comparison on the Friedman test statistics with Bonferroni correction).

Overall, the strength and effective frequency range of rate-place coding showed a moderate degradation as sound level increased. Although the performance of human listeners in F0 discrimination for HCT with resolved harmonics degrades somewhat with increasing stimulus level (Bernstein and Oxenham, 2006), the degradation does not occur until fairly high levels (> 70 dB SPL). It is unclear to what degree such degradation in behavioral performance can be accounted by the dependence of the IC rate-place code on sound level.

### Spectral receptive field model suggests a role for inhibition in rate place coding

In many rate-place coding neurons, e.g. Neuron A (Figure 4B), the trough firing rates between peaks associated with resolved harmonics were below the neuron’s spontaneous firing rate, and sometimes even reached zero, suggesting inhibition or suppression at these F0s. Such response patterns contrast with the rate-place profiles measured in the auditory nerve of anesthetized cats (Cedolin and Delgutte, 2005) where the trough firing rates were always above spontaneous rates. To explore possible mechanisms of the suppressed IC response, we fitted two spectral receptive field models to the rate-place profiles of neurons demonstrating rate-place coding: a Gaussian function with a single excitatory band and a Difference of Gaussian (DoG) function with both excitatory and inhibitory bands centered at the same frequency. When the inhibition is broader than the excitation in the DoG model, the net receptive field has a center excitatory band flanked by two inhibitory sidebands (see Methods). For both models, the parameters (center frequency, bandwidths, and, for the DoG, relative amplitude of excitatory and inhibitory components) were fitted to minimize the distance between predicted and measured rate profiles.

Figure 7A shows the Gaussian (green) and DoG (purple) fits to the rate-place profile of Neuron A at 30 dB (blue with error bars). Both models produced a rate-place profile with multiple peaks at integer harmonic numbers. However, the DoG model was better at capturing the peak amplitudes: high rates for the first three peaks followed by a much lower rate for the 4^th^ peak, and an almost indistinguishable 5^th^ peak. In contrast, the Gaussian model produced similar peak firing rates for NH from 1 to 5, thereby underestimating the amplitudes of first three peaks and overestimating the height of the 5^th^ peak compared to the neural data. The best fitting DoG and Gaussian models are shown in Figure 7B along with key model parameters. For both models, the best fitting center frequencies were close to the adjusted CF obtained by the Fourier method. The finding that the DoG model fits better than the Gaussian model in this neuron suggests a role for inhibition in shaping the rate-place profile. Inhibition is also apparent in the neuron’s FRA (Figure 4A), where the firing rate fell below spontaneous rate in frequency bands (dark blue) on either side of both center excitatory zones.

**Figure 7.**
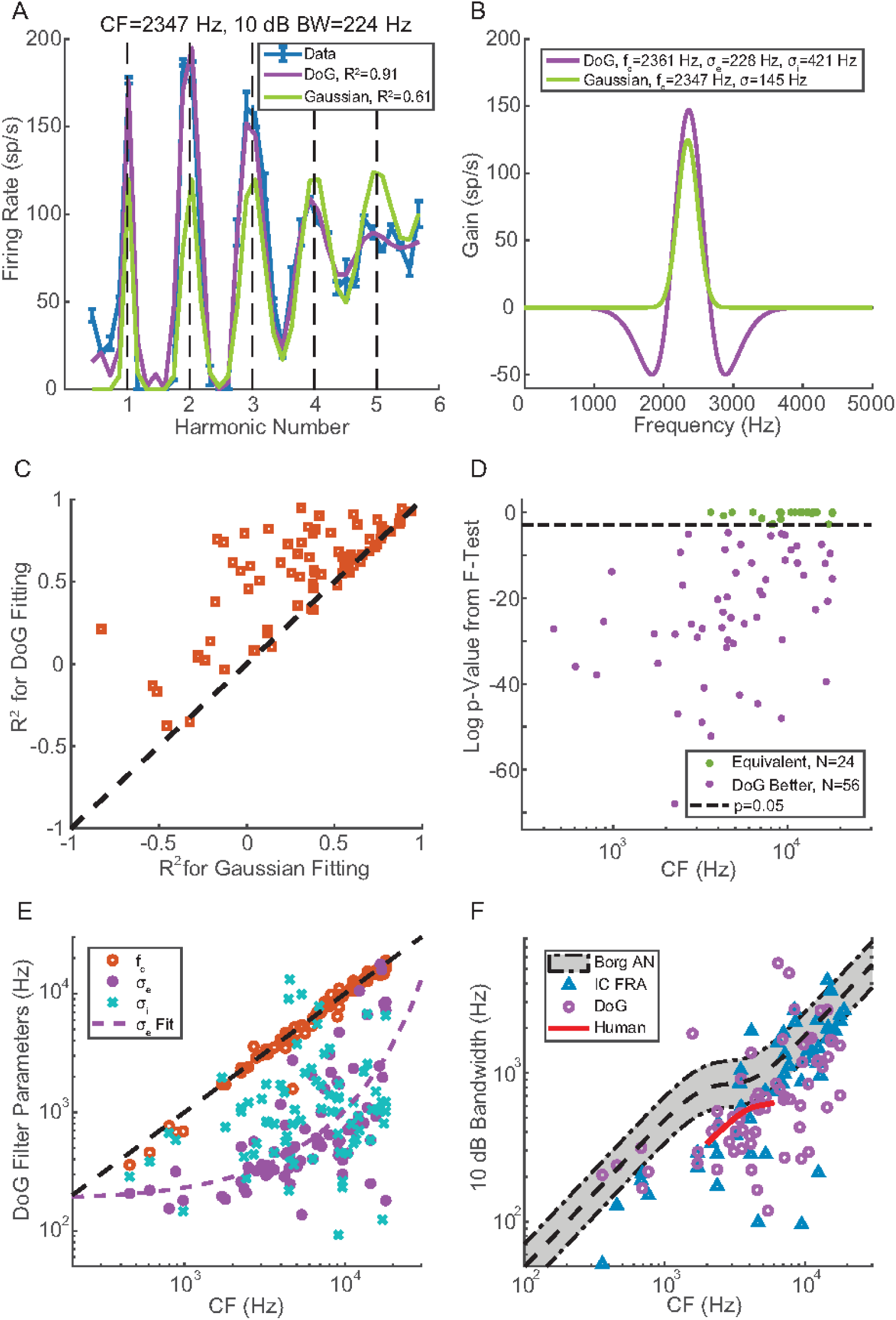
Receptive field model for harmonic complex tone responses. **(A)** Neural (blue with error bar) and fitted (purple, DoG model; green, Gaussian model) rate-place profiles of Neuron A at 30 dB SPL. 10 dB bandwidth was calculated from FRA at 30 dB SPL. **(B)** Morphology and key parameters of the optimal DoG (purple) and Gaussian (green) weighting functions. **(C)** Comparison of R^2^ values using the DoG model (y axis) versus the Gaussian model (x axis) for individual neurons (N=80). **(D)** F test statistics to compare the goodness-of-fit from the two models. Dashed line indicate p=0.05, the criterion for statistical significance. **(E)** CF (orange circle), excitatory bandwidth (purple dots), and inhibitory bandwidth (blue “x”) of the optimized DoG model as a function of measured CF in individual neurons. Purple dashed curve indicate exponential fit of the σ_e_ vs. CF relationship. **(F)** 10 dB bandwidth estimated from Borg et al. AN data (grey shaded area, 95% boundary of original data points), FRA of IC neurons showing rate-place coding (N=53, blue triangle), DoG fitting of IC neurons with rate-place coding (N=68, purple circle), and human periphery (short red curve) as functions of CF. Black dashed line at the center of the shaded area indicates a custom fitting of Borg et al. data.

In Figure 7C, the goodness of fit for the two models is compared in each neuron using the adjusted R-squared metric. Most neurons (62/80) showed a higher R-squared for the DoG model than for the Gaussian model. For 70% of neurons (56/80), the DoG fit yielded R^2^>0.5. Further examination of neurons with R^2^<0.5 in DoG fits reveals that the rate-place profiles of these neurons were either irregular (e.g. containing minor peaks between integer harmonic numbers), or showed a dramatic difference in the peak firing rates at different harmonic numbers (e.g. very high peak firing rates at NH=1 and 2, but much lower rates at NH=3, 4,…). The mechanism yielding such rate-place profiles is not clear.

To further verify the benefit of adding inhibition to the model, we ran F-tests in each neuron to compare the goodness of fit for the two models. As shown in Figure 7D, the F test p value was smaller than 0.05 for the majority of neurons (N=62), indicating that the DoG model fit the rate profiles significantly better than the Gaussian model. Equally good fits from the two models was only observed in neurons with CF >1500 Hz, but this may be just due to the smaller number of low-CF neurons in our sample.

The center frequency (f_c_, orange circle), excitatory bandwidth (σ_e_, purple dot), and inhibitory bandwidth (σ_i_, blue “x”) of the best-fitting DoG model are shown for each neuron as functions of the neuron’s adjusted CF in Figure 7E. The f_c_ values were all distributed along the identity line (black dashed), as expected. The dependence of the excitatory bandwidth of the DoG filter on CF (both in Hz) could be approximated by an exponential function shown as the purple dashed curve. In contrast, the inhibitory bandwidths were very scattered and a clear CF dependence could not be identified.

In effect, the inhibition in the DoG model sharpens the central excitation band. Figure 7F shows the relationship between CF and 10 dB bandwidth of IC neurons calculated from the central excitatory band of the DoG model (purple circles, N=68, fits with negative R^2^ or bandwidth >8000 Hz excluded). The bandwidths showed an increasing trend with increasing CF, but the data points were very scattered. A similar trend can be discerned for 10 dB bandwidths calculated from the pure-tone FRA at the same sound level as the HCTs (blue triangles, N=53, only place coding neurons included). There was no significant difference between the two measures of bandwidths from the same neuron across the population (p=0.49, two sided Wilcoxon signed-rank test).

To compare the IC bandwidths to those in the periphery, we fit a sigmoidal function to the Q_10_-CF relationship measured in rabbit auditory nerve by Borg et al. (1988), and calculated the 10 dB bandwidths from the fitted curve (black dashed line). The grey shaded area encompasses 95% of the data points for rabbit AN 10 dB bandwidths. A majority of both FRA and DoG 10 dB bandwidths for IC neurons lay below the lower bound of AN bandwidths. Both IC bandwidths were significantly smaller than the AN bandwidths (p<10^-6^ for DoG, p<10^-4^ for FRA, single-sided Wilcoxon signed-rank tests), suggesting a sharpening of frequency tuning in the auditory midbrain, at least for place-coding neurons.

### IC neural responses to inharmonic tones

When all harmonics of an HCT containing resolved harmonics are shifted in frequency by the same amount, the perceived pitch of the resulting inharmonic complex tone shifts in the same direction as the frequency shift (Schouten et al., 1962), although the temporal envelope repetition rate is unchanged (Figure 1). To shed light on mechanisms of pitch shift perception and to test whether IC neurons are sensitive to the harmonicity of complex tones, we measured the response to harmonic and inharmonic tones in 25 neurons.

Figure 8A (solid lines) shows the firing rate of Neuron C (CF=3.2 kHz) in response to complex tones with and without frequency shifts plotted against the “pseudo harmonic number” pNH=CF/FS (where FS is the frequency separation between adjacent frequency components, equal to F0 in the harmonic case). In the 0% shift (harmonic) condition, the neuron showed distinct peaks at NH=1, 2, 3. With inharmonic tones, the neuron’s profiles were similar to those in the harmonic condition with respect to peak firing rates and numbers of peaks, but were shifted in the same direction as the frequency components. Peaks in the inharmonic profile roughly occurred when pNH=integer + shift, consistent with the prediction that firing rate is maximum when a component of the shifted tone aligns with the CF (see Method).

**Figure 8.**
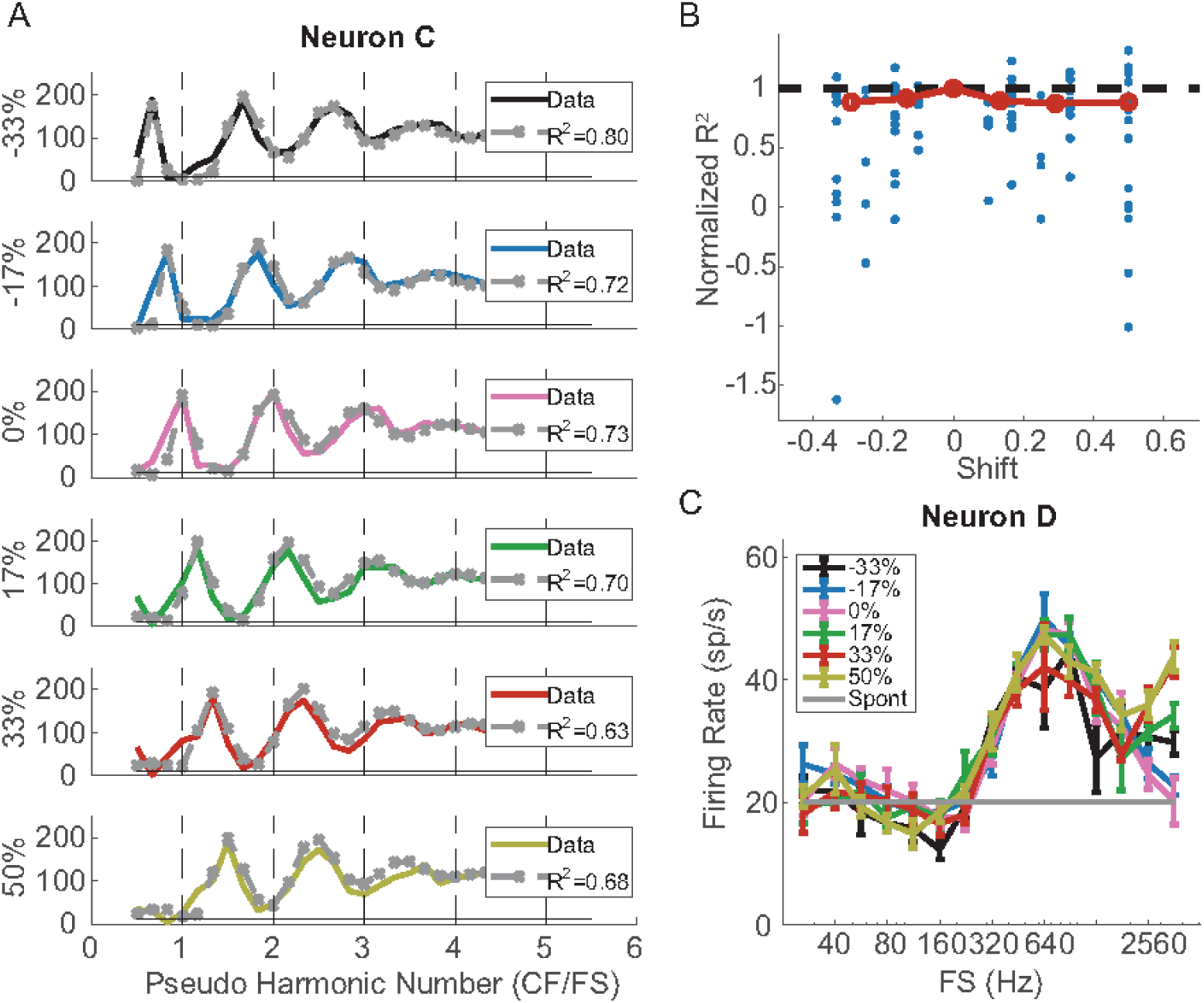
Neural and model responses to inharmonic complex tones. **(A)** Rate-place profiles of Neuron C in response to harmonic and inharmonic tones at 30 dB per component (solid lines). A DoG model was fit to the response to 0% shift then used to predict responses to non-zero shifts (grey dashed lines with “x”). **(B)** Normalized R^2^ values of DoG fitting to rate-place profiles at different proportions of frequency shift in individual neurons (blue dots) and their medians (red circles). **(C)** Rate-FS profile of Neuron D at different amounts of frequency shift.

To further verify that the neuron’s responses to the frequency-shifted tones were not dependent on harmonicity, we first fit a DoG model to the rate-place profile for 0% shift, and then used the same model parameters to predict the rate profiles in response to inharmonic tones (Figure 3.8A, dashed lines) using the shifted power spectra as inputs to the model. The DoG model predictions were very close to the neuron’s firing rates in all conditions. Adjusted R^2^ values indicated similar goodness-of-fit across the different amounts of shift. Among the 25 neurons tested with inharmonic tones, five either had negative R^2^ values in the 0% shift condition, or showed weak rate place coding. For the remaining 20 neurons, we computed the normalized R^2^ as a function of frequency shift (Figure 8B), where the R^2^ values at non-zero shifts were normalized by the value at 0% shift in the same neuron. Although the median normalized R^2^ slightly decreased with increasing absolute shift, the trend was not statistically significant (Kendall’s tau =0.157, p=0.8304).

It is worth noting that because the shifted rate-place profiles were dependent on a neuron’s ability to resolve frequency components, not all neurons showed an effect of frequency shift. For example, Neuron D (CF=9.05 kHz) in Figure 8C did not show peaks in firing rate for resolved harmonics for either harmonic or inharmonic tones. Its rate profiles for inharmonic tones were almost identical to the profile in the harmonic case, with a broad peak at 640 Hz that was not a subharmonic of the CF. By manipulating the phase relationships among the harmonics to dissociate F0 from envelope repetition rate in the stimulus waveform (Su and Delgutte, 2019), we ascertained that this neuron was sensitive to envelope repetition rate rather than F0 *per se* (not shown). Such envelope sensitivity is consistent with the lack of an effect of frequency shift on rate profiles because the envelope repetition rate equals 1/Fs regardless of the amount of shift (Figure 1B).

### Partial evidence for “harmonic template neurons” in IC

In a recent study, Feng and Wang (2017) identified a class of “harmonic template neurons” (HTN) in the auditory cortex of awake marmoset monkeys that were defined by two properties: 1) Facilitation: the firing rate in response to an HCT at the “Best F0” (BF0, which evokes the maximum firing rate to HCTs) was at least 100% higher than the rate for a pure tone at the best frequency (BF), and 2) Shift periodicity: in response to inharmonic tones created by shifting all the frequency component of a HCT by the same amount, the firing rate showed a quasi-periodic pattern as a function of the amount of shift, with maxima at integer multiples of BF0. The above results with inharmonic tones show that the shift-periodicity property holds for rate-place coding IC neurons. Therefore, we focused on testing the facilitation property of HTN by comparing the firing rates produced by pure tones and HCT as a function of frequency. Although Feng and Wang tested only one stimulus level in each neuron, we analyzed responses at three sound levels to assess whether the “harmonic template” properties are dependent on intensity.

To test the facilitation property of HTN, we measured responses to both pure tones near the CF and HCT stimuli at the same stimulus level per harmonic for 37 neurons, 22 of which showed rate-place coding of resolved harmonics. The pure tone FRA and rate profiles for pure and complex tones of Neuron E are shown in Figure 9A and 9B, respectively. This low-CF neuron (980 Hz) could resolve the first two harmonics at all three stimulus levels. The Best F0 producing the highest firing rate for HCT was equal to the CF at 45 and 60 dB SPL/component but to CF/2 at 30 dB SPL/component, indicating the neuron was more responsive to the second harmonic than the fundamental at this SPL. Such preference for a particular harmonic other than the fundamental was common in our neuronal sample. Figure 9C shows the distribution of preferred harmonics for all neurons demonstrating rate-place coding (N=80). For all three sound levels, more than one quarter of the neurons preferred the 2^nd^ harmonic over the first, and some responded strongest to even higher harmonics up to the 8^th^. The preferred harmonic shifted slightly towards lower values as sound level increased, but the trend was not statistically significant (chi-square test, p=0.28). The distribution of preferred harmonics for rate-place-coding IC neurons qualitatively resembles the distribution for cortical harmonic template neurons (Figure 4A in Feng and Wang (2017)), but the mode of the distribution is at the second harmonic for cortical neurons as opposed to the fundamental for IC neurons.

**Figure 9.**
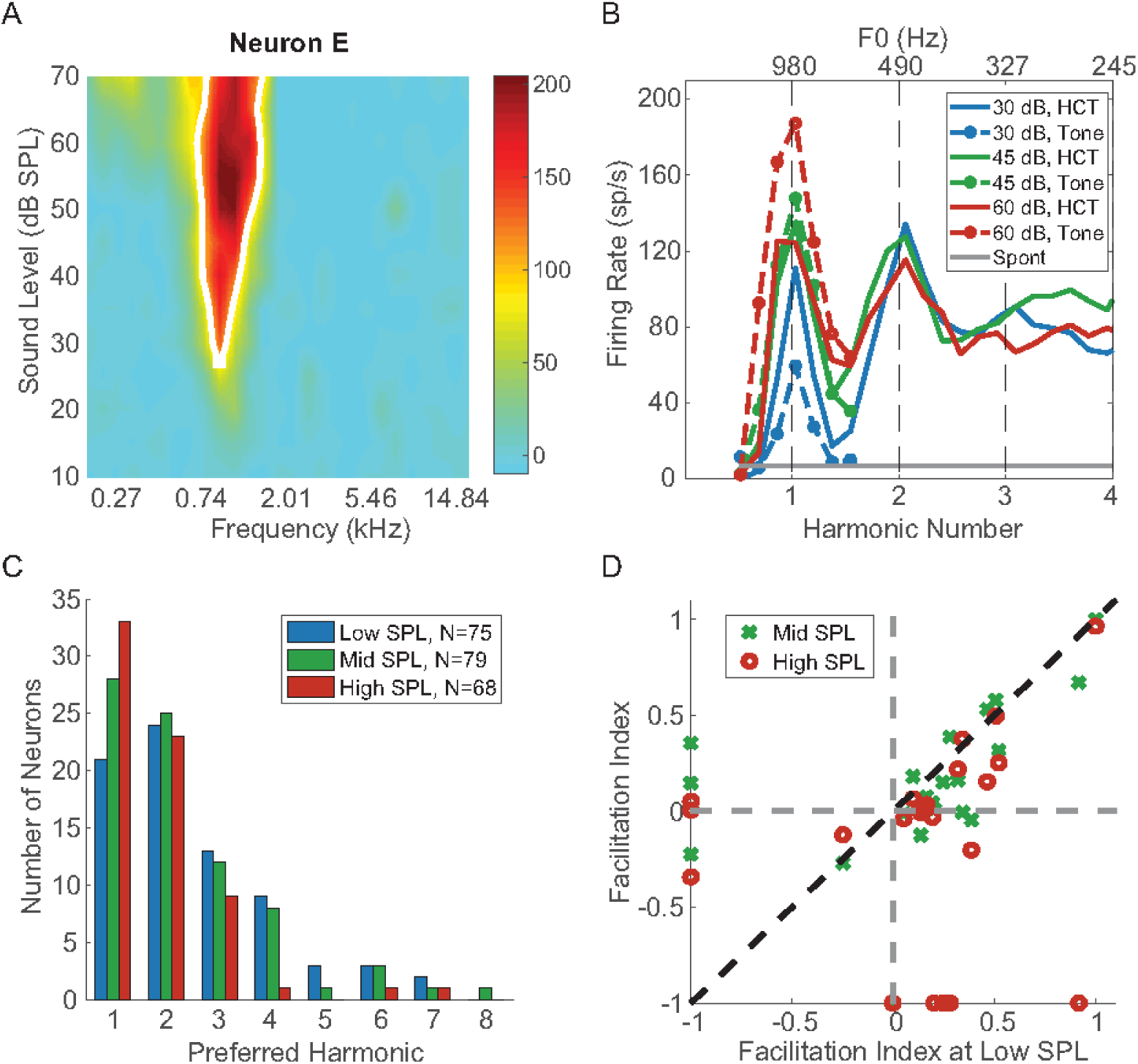
Evidence of harmonic templates in IC. **(A)** Pure tone FRA of Neuron E. **(B)** Rate profile of Neuron E in response to pure tone and HCT at different sound levels. The original measurement used HCT with harmonic number up to 6.5, and is truncated here to show detail. **(C)** Distribution of preferred harmonic across all neurons demonstrating rate-place coding (N=80). **(D)** Facilitation indices of neurons measured with interleaved pure tone and HCT, and showing rate-place coding (N=22).

Feng and Wang (2017) defined a “facilitation index” (FI) to quantify the enhancement of firing rate for HCT relative to pure tones: FI = (FR_BF0_ – FR_BF_)/(FR_BF0_ + FR_BF_). The neuron of Figure 9B shows facilitation at 30 dB SPL (FI = 0.3), but not at 45 and 60 dB, where the response to HCT is actually suppressed relative to the CF tone response. This occurred because the firing rate increased with stimulus level for the pure tone at CF, but stayed nearly constant across levels for the HCT at BF0. Figure 9D shows a scatter plot of FI at a mid and high levels against FI at the low level for the 22 rate-place coding neurons tested with both pure tone and HCT stimuli. Some neurons did not demonstrate rate-place coding at all three sound levels, and therefore did not have a Best F0 at some sound levels. Such neurons are plotted along either the x axis (neurons with a best F0 at the low SPL but not at mid or high SPLs) or the y axis (neurons with a best F0 at mid or high SPL but not at the low SPL). By definition, FI>0 indicates the neuron’s firing rate was facilitated for a HCT at BF0 compared to a pure tone at CF, while FI<0 indicates suppression for HCT. Among the 22 neurons, 17 showed facilitation at the lowest sound level, but only 5 showed suppression. The number of facilitated neurons decreased to 15 at the medium sound level, and 11 at the highest sound level. For a given neuron, FI was usually maximum at the low SPL, as data for most neurons lay under the identity line in Figure 9D. While facilitation was observed in many neurons, only 9 neurons reached the FI>0.33 criterion of Feng and Wang (2017) for HTN, which means the firing rate for a HCT at BF0 was at least 100% higher than the rate for a pure tone at CF. Overall, facilitation of HCT responses relative to pure tone responses in IC neurons was not as strong as for cortical HTNs and its prevalence was dependent on stimulus intensity. However, because we only tested IC neurons that had an identifiable pure tone CF, while some HTNs in the marmoset cortex did not respond to pure tones, we may have missed possible HTN-like neurons that responded poorly to pure tones. For the few neurons with FI<0, the FI values were fairly close to 0, meaning the amount of suppression for HCT was only modest. The dependence of suppression on sound level cannot be reliably assessed due to the small sample size.

## DISCUSSION

Using single-neuron recordings from the IC of unanesthetized rabbits in response to harmonic and inharmonic complex tones, we characterized rate-place coding of resolved frequency components, which was observed mainly for F0>800 Hz. Many IC neurons could resolve the first 3-5 harmonics, and this rate-place code was moderately dependent on sound level and not sensitive to harmonicity. Using spectral receptive field models, we found indirect evidence that inhibition may enhance the IC rate-place code. Some IC neurons had some properties of cortical harmonic template neurons in that they demonstrated facilitation to HCT compared to pure tones at CF or responded most strongly to higher harmonics rather than to the fundamental.

### Rate-place coding along the auditory pathway

Rate-place coding of resolved harmonics in HCT has been described in AN fibers (Cedolin and Delgutte 2005, 2010) in anesthetized cats and single- and multi-units in A1 of awake macaques (Schwarz and Tomlinson, 1990; Fishman et al., 2013). In the AN, rate-place coding of resolved harmonics is observed in neurons with CFs above 400 Hz, and can encode F0s above 400-500 Hz (Cedolin and Delgutte, 2005). However, the AN rate-place code degrades rapidly with increasing sound level. Although IC rate-place coding appears to be more robust across sound levels than its AN counterpart, a quantitative comparison with the data of Cedolin and Delgutte (2015, 2010) cannot be made due to differences in both stimuli (different NH ranges) and preparation (unanesthetized rabbit vs. anesthetized cat).

In both cortical studies (Schwarz and Tomlinson, 1990; Fishman et al., 2013), rate-place coding was also more prevalent in neurons with higher CFs (>300-400 Hz), and was only effective for F0s above ∼80 Hz. Schwarz and Tomlinson (1990) observed peaks in firing rates at resolved harmonics over a 40 dB range of SPLs, and Fishman et al. (2013) stated that the rate-place code was prominent at 60 dB SPL per harmonic. Thus the cortical representation was more robust against sound levels compared to the peripheral code.

In the current study, 41% of IC neurons demonstrated rate-place coding of resolved harmonics. The CFs of these neurons were all above 600 Hz, and the lowest resolved F0 was ∼300 Hz. Comparing to the single-unit results from cat AN (Cedolin and Delgutte, 2005) and macaque A1 (Schwarz and Tomlinson, 1990), the proportion of rate-place coding neurons shows a decreasing trend along the ascending auditory pathway, while the CF ranges of these neurons are comparable at different stages. In addition, IC rate-place coding was observed up to 70 dB SPL per harmonic, and was only moderately dependent on stimulus level. This evidence suggests a role for the IC in transforming the rate-place code between the periphery and the auditory cortex. However, comparisons are made difficult by differences in methodology and also known differences between cats, rabbits and macaques with respect to cochlear frequency selectivity (Liberman, 1978; Borg et al., 1988; Joris et al., 2011).

### Role of Inhibition in IC

Our experimental and modeling results suggest that inhibition may play a role in shaping rate-place coding in IC neurons. Specifically, we found that the 10-dB bandwidths of rate-place coding IC neurons were narrower than those of AN fibers (Figure 7F). A sharpening of frequency tuning relative to the AN has been observed for some classes of IC neurons with pure-tone stimulation (Ramachandran et al., 1999; Palmer et al., 2013). A recent study in mice also suggested a role of inhibition in shaping the pure-tone frequency selectivity of IC neurons (Lee et al., 2019). Our observations are consistent with studies showing a central excitatory area flanked by inhibitory bands in spectral or spectro-temporal receptive fields of IC neurons measured with broadband stimuli (Qiu et al., 2003; Lesica and Grothe, 2008; Yu and Young, 2013). However, using binaural broadband noise stimuli, McLaughlin et al. (2007) found no difference between AN and IC bandwidths in low-CF neurons sensitive to interaural time differences.

Another possible role for inhibition is to make the neural response invariant to stimulus intensity, which is highly relevant for pitch and timbre perception. Lesica and Grothe (2008) found that the STRF of IC neurons often displayed more inhibitory regions at high SPLs compared to low-SPLs. They suggested that the increase in inhibition at high levels can be attributed to the activation of high-threshold inhibitory inputs, which is in turn supported by intracellular studies that showed higher response thresholds for inhibitory compared to excitatory inputs (Covey et al., 1996; Xie et al., 2007). Thus, our finding that IC rate-place coding is fairly robust against sound level, together with previous studies showing level-invariant tuning in the primary auditory cortex (Sutter, 2000; Sadagopan and Wang, 2008), suggest that the neural representation of auditory stimuli may progressively become invariant to intensity by accumulating intensity-dependent inhibition along the ascending pathway.

### Harmonic templates for pitch extraction

It has long been known that models of pitch processing incorporating internal harmonic templates can account for a wide variety of pitch phenomena for stimuli with resolved harmonics, including the pitch shift of inharmonic complex tones (Goldstein, 1973; Wightman, 1973; Terhardt, 1974; Gerson and Goldstein, 1978; Cohen et al., 1995). Shamma and Klein (2000) proposed a biologically-plausible model for the emergence of harmonic templates based on across-frequency coincidence detection. Because the operation of coincidence detectors requires precise phase locking, the harmonic templates in this model must emerge early on in the auditory pathway, and not beyond the IC (Shamma and Dutta, 2019) where phase locking degrades sharply (Joris et al. 2004). So far neurons with properties of harmonic templates have only been reported in auditory cortex (Feng and Wang, 2017).

We explored whether harmonic templates might already arise in the midbrain by comparing IC neural response properties to the HTNs identified in marmoset auditory cortex. Specifically, we tested whether rate-place coding IC neurons meet the facilitation property of HTNs according to Feng and Wang (2017). (The other defining property – periodicity in response to shifted tones - necessarily holds for rate-place coding neurons.) A majority of IC neurons showed facilitation in response to harmonic complex tones compared to pure tones, but to a lesser degree than cortical HTNs because: 1) only a few neurons showed >100% facilitation, and 2) fewer IC neurons responded maximally to high harmonics compared to cortical HTNs. Both of these properties were dependent on sound level. We did not test neural responses to spectral jitter of the harmonics, which provided some of the strongest evidence for harmonic templates in the Feng and Wang study.

Overall, our results suggest that harmonic templates may arise gradually along the auditory pathway rather than abruptly in auditory cortex.

### Implication for pitch perception

The lowest F0 for which we found rate-place coding of resolved harmonics in rabbit IC (∼300 Hz) lies at the upper range of adult human voice (∼80-320 Hz) (Matteson et al., 2013), and the most effective F0 range for IC rate-place coding (>3000 Hz, Fig. 5D) lies mostly above the upper limit of musical pitch around 5 kHz (Semal and Demany, 1990; Plack and Oxenham, 2005a). Such differences can be partly attributed to broader cochlear tuning in rabbit, as shown in Figure 7F by the comparison of 10-dB bandwidths of rabbit ANF (Borg et al., 1988) with estimates of human AN bandwidths from compound action potentials (Verschooten et al., 2018). Extrapolating the trends in human AN bandwidths to lower CFs, the bandwidths of neurons with CFs in the first formant region (200-800 Hz) may be narrow enough to resolve harmonics of F0s in the range of male voices, and this would also be true for human IC neurons if the sharpening observed in the rabbit IC also occurs in humans The few available studies on rabbit phonation suggest F0s in the range of 500-1200 Hz (Swanson et al., 2010; Mills et al., 2017; Döllinger et al., 2018), which is at the low end of the effective range for rate-place coding by IC neurons.

Human psychophysical experiments using HCTs with identical power spectra but different temporal envelope repetition rates (ERR) (Krumbholz et al., 2000; Pressnitzer et al., 2001) showed that, at very low frequencies (<50 Hz), the pitch of harmonic complex tones is dependent on ERR rather than F0. Therefore, the lower limit of pitch perception is likely not conveyed by resolved spectral components, but by temporal phase locking to the ERR. In a companion paper (Su and Delgutte, 2019), we showed that a temporal code for ERR was available in the IC up to 900 Hz, and a rate tuning to ERR was observed between 56 and 1600 Hz. Therefore, the three neural codes available in IC are effective in complementary frequency ranges, and together cover a wide range of F0.

Our findings on IC rate-place coding shed light on the transformation of neural representations of resolved harmonics along the auditory neuraxis, and suggest how a higher processing center could extract pitch from resolved frequency components. To understand the perceptual relevance of the IC neural representations, it is necessary to test perception of complex tones in rabbits with behavioral methods (Delgutte et al., 2018).

## Acknowledgement

This work is supported by NIH grant R01 DC 002258. We thank Yoojin Chung for help with experimental procedures, Ken Hancock for technical support, Oded Barzelay for providing the algorithm for estimating characteristic frequency, Kameron Clayton for providing the fitting of rabbit auditory nerve bandwidth, Camille Shaw, Alice Gelman, and Joseph Wagner for assisting with surgeries.

